# Differential dysregulation of β-TrCP1 and -2 by HIV-1 Vpu leads to inhibition of canonical and non-canonical NF-κB pathways in infected cells

**DOI:** 10.1101/2022.11.30.518636

**Authors:** Suzanne Pickering, Jonathan Sumner, Claire Kerridge, Stuart Neil

## Abstract

The HIV-1 Vpu protein is expressed late in the virus lifecycle to promote infectious virus production and avoid innate and adaptive immunity. This includes the inhibition of the NF-κB pathway which, when activated, leads to the induction of inflammatory responses and the promotion of antiviral immunity. Here we demonstrate that Vpu can inhibit both canonical and non-canonical NF-κB pathways, through the direct inhibition of the F-box protein β-TrCP, the substrate recognition portion of the Skp1-Cul1-F-box (SCF)^β-TrCP^ ubiquitin ligase complex. There are two paralogues of β-TrCP (β-TrCP1/BTRC and β-TrCP2/FBXW11), encoded on different chromosomes, which appear to be functionally redundant. Vpu, however, is one of the few β-TrCP substrates to differentiate between the two paralogues. We have found that patient-derived alleles of Vpu, unlike those from lab-adapted viruses, trigger the degradation of β-TrCP1 while co-opting its paralogue β-TrCP2 for the degradation of cellular targets of Vpu, such as CD4. The potency of this dual inhibition correlates with stabilisation of the classical IκBα and the phosphorylated precursors of the mature DNA-binding subunits of canonical and non-canonical NF-κB pathways, p105/NFκB1 and p100/NFκB2, in HIV-1 infected CD4+ T cells. Both precursors act as alternative IκBs in their own right, thus reinforcing NF-κB inhibition at steady state and upon activation with either selective canonical or non-canonical NF-κB stimuli. These data reveal the complex regulation of NF-κB late in the viral replication cycle, with consequences for both the pathogenesis of HIV/AIDS and the use of NF-κB-modulating drugs in HIV cure strategies.

## Introduction

The NF-κB family of inducible transcription factors plays a fundamental role in regulating mammalian immune responses, including the induction of a pro-inflammatory state following the sensing of virus invasion. Viruses, in turn, often deploy multiple strategies to thwart sensing pathways before signalling cascades can be fulfilled. As is the case with many viruses, the interplay between HIV-1 and the NF-κB pathway is complex. The virus contains NF-κB response elements in its long terminal repeat promoter, and thus relies on NF-κB activation for the transcription of its genes [1], while also encoding inhibitory factors at different stages of the viral life-cycle – specifically, the accessory proteins Vpr and Vpu. Vpr is packaged into the virus particle and modulates the cellular environment early in infection [2, 3], while Vpu is expressed late in the virus lifecycle in tandem with the envelope protein and performs multiple functions to achieve optimal cellular conditions for virus production [4-13].

The NF-κB transcription factor family consists of NF-κB1 p50, NF-κB2 p52, p65 (RelA), RelB and c-Rel, that associate in homo-or heterodimers and are activated by canonical and non-canonical pathways (**Figure 1a** ; [14, 15]). The canonical pathway is responsive, rapid and transient, responding to stimuli such as pattern recognition receptors (PRRs), inflammatory cytokines (including TNFα and IL-1β), and antigen receptors to mediate essential roles in innate and adaptive immunity [16]. In the paradigm canonical pathway, NF-κB dimers, most commonly p65/p50, are held inactive in the cytoplasm by inhibitors of κB (IκB), including IκBα and the precursor IκBs p105 (also called IκBγ) and p100 (also called IκBδ). Stimulation of the pathway activates the IκB kinase (IKK) complex, which phosphorylates IκBs, leading to their ubiquitination and proteasomal degradation (**Figure 1a**). This releases the NF-κB transcription factor for translocation to the nucleus and transcription of NF-κB-dependent target genes containing NF-κB-dependent response elements (GGGRNNYYCC) in their promoters [16].

**Figure 1.**
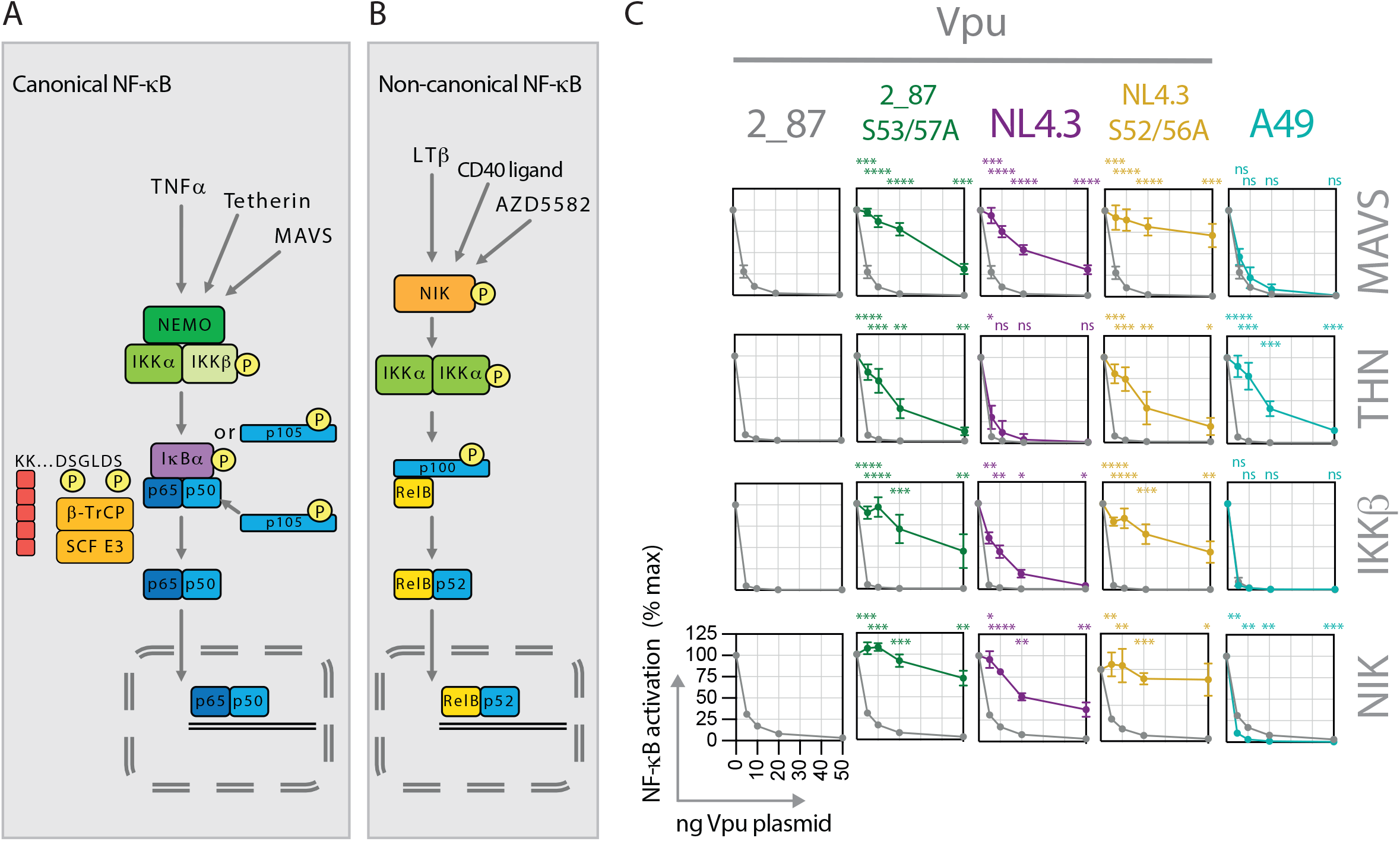
Vpu inhibits both the canonical and the non-canonical NF-κB pathways. **(A)** Graphical representation of the canonical NF-κB pathway, detailing events downstream of the activation of the IKK complex. Stimuli such as TNFα, tetherin activation (through the retention of budding virus particles) or MAVS activation (following upstream sensing of viral RNA) trigger signalling cascades that converge at the activation of the IKK complex. IKKβ phosphorylates inhibitors of NF-κB, most commonly IκBα (but also p105 and p100), on dual serine residues 32 and 36 in the degron sequence SGLDS, leading to recognition by the β-TrCP substrate adaptor portion of an E3 cullin-RING ligase (SCF^β-TrCP^). Ubiquitination of IκBα on lysine residues by SCF^β-TrCP^ triggers proteasomal degradation, releasing the NF-κB transcription factor (in this example the p65/p50 heterodimer), which translocates to the nucleus and activates the expression of NF-κB-dependent genes. P105 also acts as a precursor for the p50 portion of the NF-κB transcription factor, and is converted to active p50 by partial proteasomal processing. **(B)** Graphical representation of the non-canonical NF-κB pathway. Stimuli such as lymphotoxin β (LTβ), CD40 ligand, or the synthetic compound AZD5582, lead to the activation of NIK, which in turn phosphorylates IKKα. Activated IKKα phosphorylates p100 on dual C-terminal serine residues, prompting its recognition by SCF^β-TrCP^, ubiquitination and partial proteasomal processing to form mature RelB/p52 dimers, able to translocate to the nucleus and activate transcription. **(C)** Transient NF-κB activation assays were performed in HEK293T cells by co-transfecting an NF-κB-dependent luciferase reporter construct (3xNF-κB pConA), a renilla luciferase control plasmid, a fixed dose of plasmid expressing an NF-κB stimulus (MAVS, tetherin, IKKβ or NIK), and an increasing dose of Vpu or A49 plasmid. 24 hours after transfection, cells were lysed and luciferase activity was determined. Results are expressed as a percentage of normalised signal recorded in the absence of Vpu or A49 (% max). Means are presented from at least 4 independent experiments, with error bars showing ± SD. The 2_87 line is shown on all graphs for comparison (grey line, grey circles), with 2_87 S3/7A in green, NL4.3 in purple, NL4.3 S2/6A in yellow and A49 in turquoise. Asterisks indicate points that differ significantly from 2_87: *p* value >0.1 (ns), <0.1 (*), <0.01 (**), <0.001 (***), <0.0001 (****).

The non-canonical pathway is activated following engagement of a subset of TNFR superfamily members such as LTβR, BAFFR and CD40 with a slower, more persistent response than the canonical pathway [14, 17]. It is required for lymphoid organ development, B cell survival and maturation and the maintenance of effector and memory T cells, and its activation is based on the processing of p100 [18]. The critical kinases in this pathway are IKKα and NIK, with NIK phosphorylating IKKα on serines in the activation loop, leading to its activation and phosphorylation of p100 (**Figure 1b**). Polyubiquitination signals the partial proteasomal processing of p100, destroying the C-terminal region and releasing it as a mature p52 molecule, most commonly in complex with RelB. The p52/RelB transcription factor is then free to translocate to the nucleus [15].

Both pathways depend on the ubiquitin-proteasome machinery at pivotal stages, including the proteasomal degradation of IκBα or other IκB family members and the partial proteasomal processing of the precursor proteins p105 and p100 to mature NF-κB subunits, p50 and p52 [16, 19]. Importantly, in their unprocessed form, both p105 and p100 act as IκBs, thus fulfilling a dual role in regulating the pathway dependent on the ubiquitin-proteasome system. The F-box protein, beta-transducin repeat-containing protein (β-TrCP), is the substrate adaptor protein of the Skp1-cullin1-F-box protein (SCF) E3 ubiquitin ligase machinery that initiates the ubiquitination of IκBs, p100 (IκBδ) and p105 (IκBγ). β-TrCP recognises a highly conserved phosphorylated motif with the consensus sequence DpSGxxpS in the N-terminal region of IκBα (DSGLDS) and the C-terminal region of p100 (DSAYGS) and p105 (DSGVETS) molecules. Phosphorylation of this motif, also called a (phospho)degron, provides a binding site for the beta-propeller repeat portion of the WD40 domain of β-TrCP [20], which links the substrate to the ubiquitin ligase machinery and targets it for proteasomal degradation, or in the case of the precursor proteins, induces partial proteasomal processing.

SCF^β-TrCP^ has numerous substrates beyond the canonical and non-canonical NF-κB pathways, including proteins involved in cell cycle regulation, autophagy and WNT signaling pathways. β-TrCP exists as two paralogues, β-TrCP1 (BTRC) and -2 (FBXW11), encoded on separate chromosomes, each with several functional isoforms [21]. The functional relevance of these different paralogues is unclear, with early mitotic inhibitor 1 (Emi1) being the only cellular target of β-TrCP to demonstrate requirement for both paralogues rather than redundancy [22].

Through viral molecular mimicry, the HIV-1 accessory protein Vpu contains an SGxxS motif akin to other targets of the SCF^β-TrCP^ (D<u>S</u>GNE<u>S</u>). Indeed, β-TrCP was first discovered through its interaction with Vpu [23], and was later ascribed its major function in the NF-κB pathway [24, 25]. The serines in the Vpu SGNES motif are highly conserved and are essential for the optimal execution of all known Vpu functions [6, 7, 12]. Phosphorylation of the serines by casein kinase II (CKII/CK2) [26, 27] creates a binding site for β-TrCP, which is then co-opted by Vpu both to inhibit the NF-κB pathway [11, 28-31] and to induce the ubiquitination and subsequent degradation of the HIV-1 receptor CD4 and the antiviral protein BST2/tetherin [23, 32-36]. Thus, unlike cellular proteins possessing SGxxS degrons that are themselves targeted for ubiquitination and degradation, Vpu acts as an adaptor protein to link the E3 ubiquitin ligase machinery to its target proteins. Interestingly, Vpu has also demonstrated preference for a single paralogue, β-TrCP2, in the counteraction of the antiviral protein Bst2/tetherin [33, 35].

Mechanistic insights into NF-κB inhibition by Vpu have been established from studies of the T cell line-adapted HIV-1 molecular clone, NL4.3 [29, 30]. These demonstrated a sequestration of β-TrCP, resulting in a block to the ubiquitination and degradation of IκB and downstream inhibition of NF-κB translocation. It has more recently been recognised that the potency of this activity has been underestimated, as the NL4.3 Vpu used for these studies has severely diminished activity compared with primary Vpus [11, 31]. Thus, a reassessment of the mechanism with primary Vpus is appropriate in order to fully understand the nature of the inhibition.

Here we investigate the interaction between HIV-1 Vpu and β-TrCP and its downstream consequences, including previously uncharacterised effects of Vpu on p105. We further document effects on the non-canonical NF-κB pathway, which may have implications for HIV latency reversal strategies. We demonstrate that inhibition of both pathways by Vpu involves the simultaneous degradation and sequestration of β-TrCP1 and -2 respectively, revealing the potential for distinct activities by these two paralogues, and illustrating the fine balance between exploiting and inhibiting the pathways.

## Results

### Vpu inhibits both the canonical and non-canonical NF-κB pathways

We first investigated the ability of Vpu to inhibit NF-κB activation induced by both the canonical and non-canonical pathways (**Figures 1a** and **1b**). We and others have previously reported that NL4.3 Vpu has suboptimal canonical NF-κB inhibitory activity compared to primary Vpus [11, 31], therefore we investigated NL4.3 alongside a highly active primary Vpu, 2_87, typical of those found in natural subtype B infections [31]. Double serine mutants of both Vpus, mutated at serines 53 and 56 for 2_87 or 52 and 56 for NL4.3 and unable to bind SCF^β-TrCP^, were included as controls (2_87 S3/7A and NL4.3 S2/6A), as was A49, a poxvirus protein with potent NF-κB inhibitory activity [37, 38]. Canonical stimuli used were: MAVS, which plays an integral role in viral RNA sensing; tetherin, which acts as a pattern recognition receptor upon inhibition of virus budding [39]; and IKKβ, part of the canonical IKK complex and pivotal to the canonical NF-κB pathway (**Figure 1a**). Inhibition of the non-canonical pathway was investigated by using NF-κB-inducing kinase (NIK) as a stimulus. Transfection of NIK leads to its activation and phosphorylation of IKKα, which in turn phosphorylates p100 at dual serine residues, leading to SCF^β-TrCP^ recognition, ubiquitination, and subsequent proteasomal processing to p52 (**Figure 1b**). All stimuli induced NF-κB activation when transfected into HEK293T cells, measured by luciferase reporter assay, to an average level of 122-(MAVS), 42-(tetherin), 110-fold (IKKβ) and 333-fold (NIK) above background. 2_87 Vpu and A49 potently inhibited NF-κB induced by all four stimuli, demonstrating that these viral antagonists can inhibit both the canonical and non-canonical NF-κB pathways (**Figure 1c**). In contrast, NL4.3 was significantly impaired across all concentrations (**Figure 1c**). Compared to their wildtype counterparts, both double serine mutants (2_87 S3/7A and NL4.3 S2/6A) were significantly defective against all stimuli, with 2_87 S3/7A showing some activity at higher concentrations. Inhibition of NF-κB induced by tetherin revealed differences between the antagonists, with the defective Vpus 2_87 S3/7A, NL4.3 and NL4.3 S2/6A all showing increased inhibitory activity, corresponding to the fact that Vpu has an independent direct antagonistic effect on tetherin, ultimately inducing its degradation. Conversely, A49, which potently inhibited NF-κB activity induced by MAVS and IKKβ, was less effective at inhibiting tetherin-mediated NF-κB stimulation. The inhibition of IKKβ-induced signaling confirms previous findings that Vpu and A49 inhibit the NF-κB pathway downstream of the activation of the IKK complex [29, 30, 37, 38]; while the inhibition of NIK implies a similar block to the non-canonical pathway, both consistent with inhibition occurring at the β-TrCP level of the pathway.

### Primary Vpu induces the degradation of β-TrCP in infected cells

Previous work on the mechanism of NF-κB inhibition by HIV Vpu has shown that β-TrCP is sequestered and stabilised [29, 30]. To investigate whether a direct effect on β-TrCP could be visualised, endogenous levels of β-TrCP were examined under conditions of natural infection by HIV-1. For this, NL4.3 viruses were engineered to express heterologous 2_87 Vpu, and mutants thereof, at endogenous levels. 48 hours following infection of cells with viruses expressing either 2_87, 2_87 S3/7A, NL4.3 or no Vpu, β-TrCP levels were examined by Western blot (**Figure 2**). Incongruous with the notion that Vpu sequesters and utilises β-TrCP for the degradation of its target proteins, we observed a significant and consistent depletion of β-TrCP in HEK293T, primary CD4+ T cells, and CD4+ Jurkat T cells infected with 2_87-expressing virus (**Figures 2a-c**). Viruses expressing NL4.3 Vpu exerted a similar but lesser effect. Degradation was rescued by treatment with proteasomal inhibitor MG132 and the NEDD8-activating enzyme (NAE) inhibitor MLN4924, which specifically blocks the activation of cullin-RING ligases (CRLs) by inhibiting their activation through neddylation (**Figure 2d** and **Supplementary Figure 1a**), indicating degradation through a proteasomal and CRL-dependent pathway. Treatment with an inhibitor of lysosomal degradation, concanamycin A, did not rescue degradation (**Supplementary Figure 1a**).

**Figure 2.**
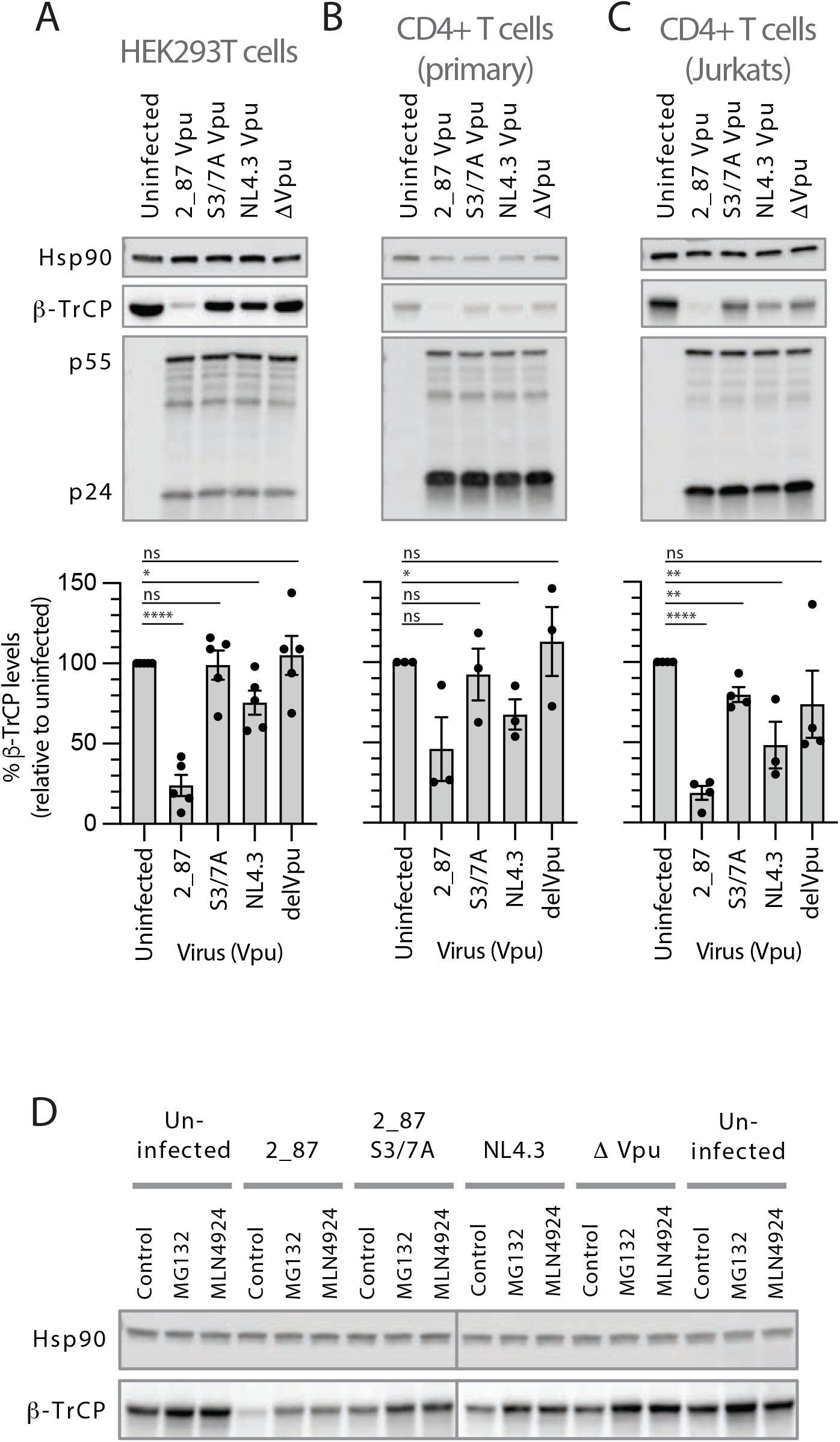
β-TrCP1 levels are significantly depleted in cells infected with virus expressing primary Vpu. Recombinant NL4.3 proviruses engineered to express either highly active 2_87 Vpu, 2_87 S3/7A Vpu, NL4.3 Vpu or no Vpu (Δ Vpu) were used to infect HEK293T cells (**A**), primary CD4+ T cells (**B**) or CD4+ Jurkat T cells (**C**) at an MOI of 5 for 48 hours. Cells were harvested and western blotted for Hsp90 (loading control), β-TrCP and HIV-1 Gag (major bands show p55 and p24). Graphs below the blots show mean β-TrCP levels from 3 to 5 independent experiments (for primary CD4+ T cells this is calculated from experiments from 3 different donors), with β-TrCP1 western blot intensities normalised first to Hsp90 for each sample, and percentages calculated relative to uninfected cells. Error bars represent ± SEM. Asterisks indicate β-TrCP levels that differ significantly from uninfected cells: *p* value >0.1 (ns), <0.1 (*), <0.01 (**), <0.001 (***), <0.0001 (****). **(D)** HEK293T cells were infected as in (**A**), but treated with proteasomal inhibitor MG132 (10 µM) or NEDD8-activating enzyme (NAE) inhibitor MLN4924 (0.1 µM) for 6 hours prior to harvest at 48 hours. Cell lysates were analysed by western blot for Hsp90 (loading control) and endogenous β-TrCP levels.

### Vpu has differential effects on β-TrCP1 and -2

The observation that Vpu leads to β-TrCP degradation is at odds with the essential role of β-TrCP in the Vpu-mediated degradation of CD4 and other cellular targets. There are two paralogues of β-TrCP – β-TrCP1 (BTRC) and β-TrCP2 (FBXW11) - encoded on separate chromosomes, each with several isoforms [21]. Previous reports have implicated β-TrCP2 in the degradation of BST2/tetherin, while β-TrCP1 was dispensable for Vpu function [33, 35]. We sought to reconcile previous reports of selective β-TrCP usage by Vpu with our observations of β-TrCP1 degradation. The lack of an antibody suitable for the detection of endogenous levels of β-TrCP2 led us to take a molecular approach. Transient expression assays were performed by co-transfecting β-TrCP1 or -2 with Vpu, in the presence or absence of active NF-κB signalling (+/-IKKβ), harvesting at 24 hours and western blotting cell lysates (**Figure 3a**). Levels of β-TrCP1 were depleted by up to 88% in the presence of 2_87 Vpu (79% in the presence of IKKβ), with S3/7A showing a modest reduction. In contrast, NL4.3 Vpu caused an almost 6-fold increase in β-TrCP1 levels (3.4-fold in the presence of IKKβ). In agreement with previous studies espousing sequestration of β-TrCP through molecular mimicry by Vpu [29, 30], we demonstrate that both 2_87 and NL4.3 robustly stabilise β-TrCP2, both in the presence and absence of stimulus (**Figure 3a** and **Supplementary Figure 1b**). Control experiments demonstrate that GFP levels remain unchanged under the same conditions (**Supplementary Figure 1b**). These data corroborate our findings in infected cells, and highlight the differential effects on β-TrCP, both in terms of β-TrCP paralogues and when comparing 2_87 and NL4.3 Vpu. Given the simultaneous reduction of β-TrCP1 and stabilisation of β-TrCP2, and the involvement of a CRL- and proteasomal-dependent degradation pathway for the former (**Figure 2d**), we hypothesised that Vpu might exploit an SCF^β-TrCP2^ for the downregulation of β-TrCP1 [40]. However, siRNA knockdown of β-TrCP2 had no apparent effect on the reduced levels of β-TrCP1 in infected cells (**Supplementary Figure 1c**). Co-immunoprecipitation experiments were performed in order to establish whether the dichotomous effects on β-TrCP were due to obvious differences in binding ability of 2_87 and NL4.3 Vpu (**Figure 3b**). In the case of β-TrCP1, MG132 was added prior to co-immunoprecipitation in order to mitigate degradation by 2_87 Vpu. Both 2_87 and NL4.3 were able to bind β-TrCP1 and -2, with no discernible difference in binding ability. As expected, the serine mutants of both Vpus were unable to bind both β-TrCPs (**Figure 3b**).

**Figure 3.**
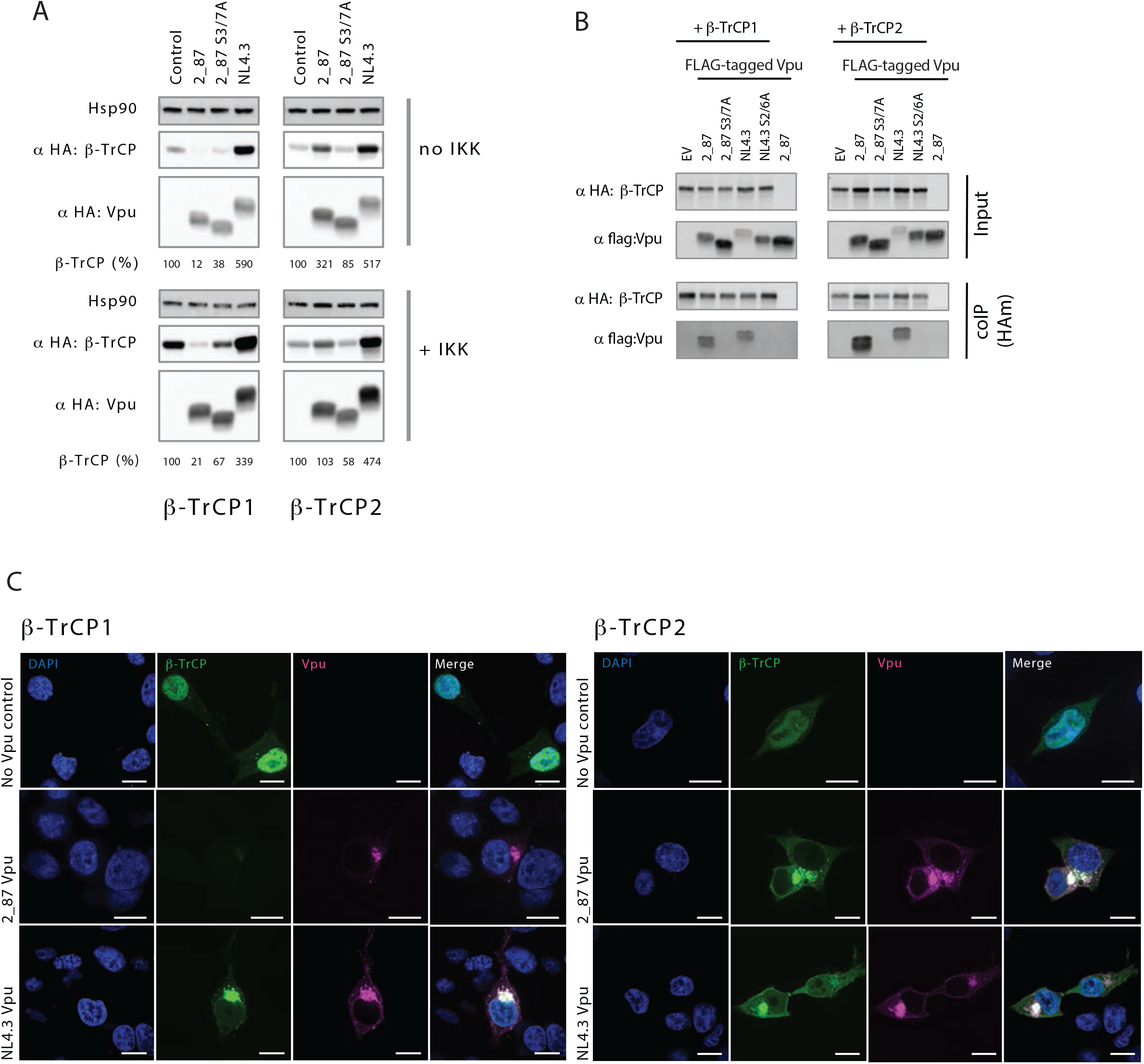
Selective degradation of β-TrCP1 by primary, but not NL4.3, Vpu. **(A)** The direct effect of Vpu on β-TrCP1 and 2 was examined by co-transfecting HEK293T cells with HA-tagged β-TrCP1 or -2 plus Vpu or empty vector control, in the presence (+IKK) or absence (no IKK; empty vector) of active signalling. 24 hours after transfection, cell lysates were harvested and analysed by western blot for HA (Vpu and β-TrCP) and Hsp90 (loading control). **(B)** 2_87 and NL4.3 Vpus were compared for their ability to bind β-TrCP1 and -2 by immunoprecipitation. Dual serine mutants of each Vpu (2_87 S3/7A and NL4.3 2/6A) were used as negative controls. HEK293T cells were co-transfected with flag-tagged Vpu or EV and HA-tagged β-TrCP or EV, and 24 hours later cells were lysed, immunoprecipitated with anti-HA antibody and analysed by western blot. **(C)** Confocal microscopy images of HEK293T cells co-transfected with GFP-tagged β-TrCP1 or - 2 (green) and HA-tagged Vpu (pink) and co-stained for DAPI (blue). Areas of colocalization appear white. Panels are single *z* slices with scale bars of 10 µm.

The binding, degradation and sequestration patterns seen in **Figures 2a-d, 3a** and **3b** were next corroborated by confocal microscopy. As shown previously [21], β-TrCP1 and -2 are found both in the nucleus and cytosol, but with predominant nuclear localisation (**Figure 3c**). In the presence of 2_87 Vpu, the previously observed contrasting effects on β-TrCP1 and -2 can be seen, with levels of β-TrCP1 severely depleted, while β-TrCP2 was dramatically re-localised to the cytosol and sequestered predominantly in perinuclear regions, consistent with typical trans-Golgi network (TGN) localisation of Vpu [41]. β-TrCP2, on the other hand, was relocalised and sequestered by both 2_87 and NL4.3 Vpu (**Figure 3c**).

### Infection with HIV-1 leads to tonic activation of NF-κB and stabilisation of p105 (NFkB1)

Such a direct and dramatic effect on β-TrCP led us to investigate the effect of primary Vpu on the β-TrCP substrates IκBα and p105, alongside all other major components downstream of the IKK complex. Cells were infected with viruses expressing the indicated Vpu and NF-κB pathway components were examined by western blot (**Figure 4a**). Total IKKβ, p65 and IκBα remained unaffected under all infection conditions, while phosphorylated p65 and IκBα were not detected. Unexpectedly, phosphorylated p105 was detected under all infection conditions but not in uninfected control cells, indicative of a vestige of ongoing NF-κB activation due to virus infection. Phosphorylated p105 was significantly increased in cells infected with 2_87 Vpu virus (**Figure 4a)**. This was also evident in both primary and Jurkat CD4+ T cells (**Figure 4b and c**). In the primary CD4+ T cells phospho-p105 was detected in all conditions, including uninfected cells, due to NF-κB activation induced by the CD3/CD28 co-stimulation conducted prior to HIV-1 infection (**Figure 4b**).

**Figure 4.**
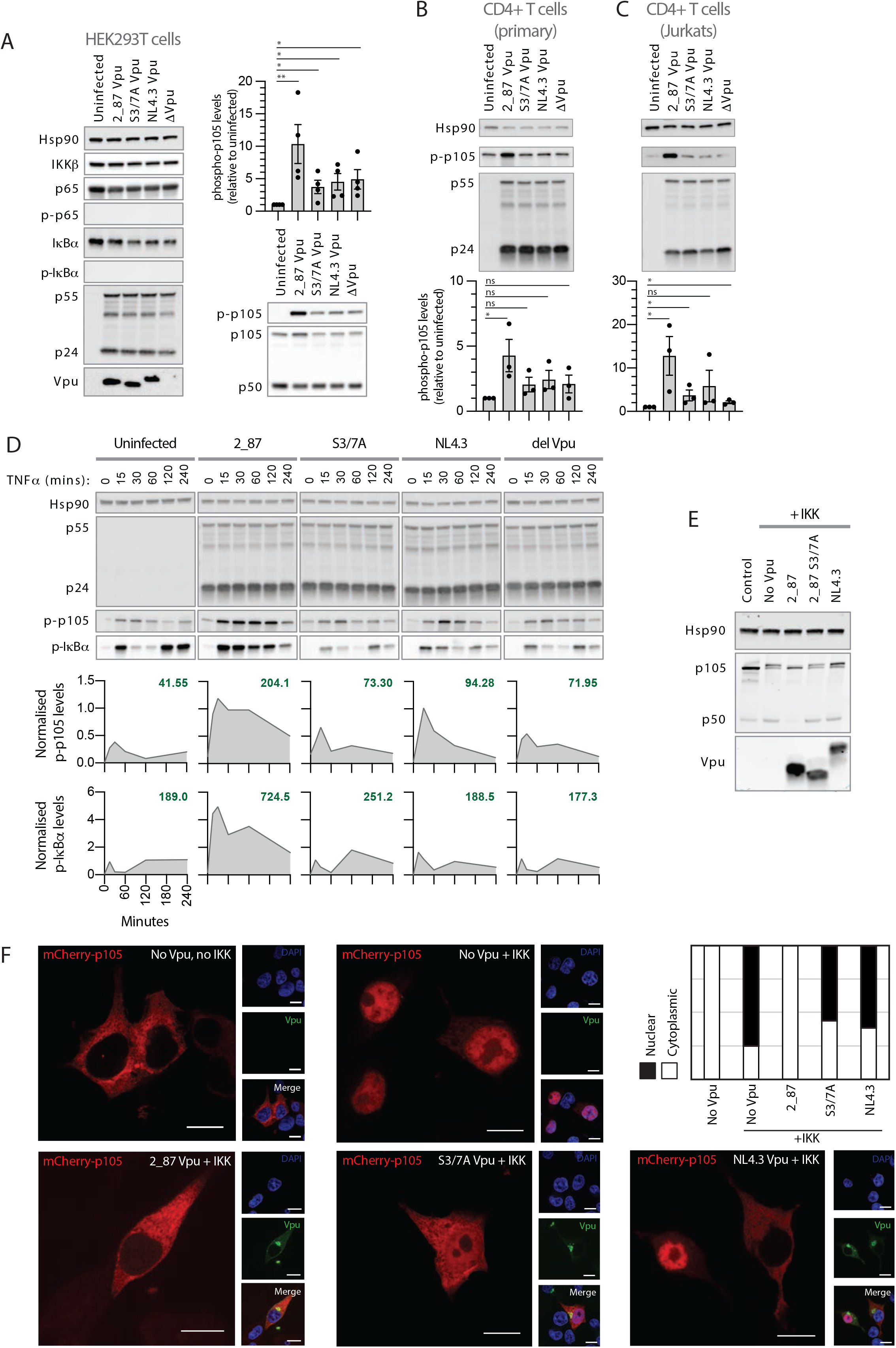
During infection and under conditions of active signalling, Vpu leads to stabilisation of p-IκBα, p-p105 and the inhibition of processing to p50 and subsequent nuclear translocation. **(A)** Components of the NF-κB complex downstream of the IKK complex (all depicted in **Figure 1A**) were examined in infected cells, in the absence of exogenous NF-κB stimulation. Recombinant NL4.3 proviruses engineered to express either highly active 2_87 Vpu, 2_87 S3/7A Vpu, NL4.3 Vpu or no Vpu (Δ Vpu), were used to infect HEK293T cells (MOI 5) for 48 hours. Cells were harvested and western blotted for Hsp90 (loading control), IKKβ, phospho-p105 (Ser932), total p105, p50, p65, phospho-p65 (Ser536), total IκBα and phospho-IκBα (Ser32/Ser36). HIV-1 Gag (p55 and p24) and Vpu were blotted as controls for infection levels. Western blots for phospho-p105 were quantified, normalised to Hsp90 levels for each lane and to the uninfected sample for each experiment, and plotted as averages of at least three separate experiments (bars). Individual data points are shown as dots. Error bars represent ± SEM. Unpaired one-tailed T tests were performed for each condition, with *p*-values indicated by asterisks: ns, not significant (p>0.05);* <0.5, **<0.05). (**B**), as for **A**, but in primary CD4+ T cells. **(C)** as for **A**, but in CD4+ Jurkat cells. **(D)** HEK293T cells were infected with viruses expressing either 2_87, 2_87 S3/7A, NL4.3 or no Vpu (Δ Vpu) at an MOI of 3. 44 hours after infection, cells were treated with 10 ng/ml TNFα, and time points were harvested at 0, 15, 30, 60, 120 and 240 minutes following treatment, resulting in a total infection duration of 48 hours. Samples were analysed by western blot for Hsp90 (loading control), HIV-1 Gag (major bands showing p55 and p24), phospho-p105, and phospho-IκBα. Band intensities for p-p105 and p-IκBα are shown below each blot, normalised to Hsp90 for each sample and to positive controls for p-105 or p-IκBα, as appropriate, per blot (not shown in the image). Numbers shown in bold green text on each graph represent the calculated area under the curve (AUC). **(E)** Transient p105 processing assays were performed by co-transfecting HA-p105, IKKβ and Vpu (2_87, 2_87 S3/7A or NL4.3) plasmids into HEK293T cells. 24 hours after transfection, cells were harvested and western blotted for HA (p105, p50 and Vpu) and Hsp90 as a loading control. **(F)** Confocal microscopy images of HEK293T cells co-transfected with mCherry-tagged p105 (red) and HA-tagged Vpu (green), in the presence or absence of active signalling (+/-IKK) and co-stained for DAPI (blue). Panels are single *z* slices with scale bars of 10 µm. Graph shows proportion of cells with nuclear p50 (white) or cytoplasmic p105/p50 (black) from 100 counted cells.

We next investigated the effect of infection under conditions of active NF-κB signalling. Infected cells were treated with TNFα, subjected to 4-hour time courses, then examined for components downstream of the IKK complex (**Figure 4d**). Typical cyclical profiles were seen for phosphorylated IκBα in the uninfected cells, with phosphorylation detected at 15 minutes, degradation at 30-60 minutes, and renewed detection of p-IκBα at 2 and 4 hours due to resynthesis of IκBα in response to NF-κB-activated transcription. P-p105 followed a similar but less pronounced profile. In cells infected with 2_87 Vpu virus, complete stabilisation of both p-p105 and p-IκBα was seen across the timecourse, with a marked increase in the detection of both p-p105 and p-IκBα alongside a loss of the degradation seen at 30-60 minutes. Cells infected with NL4.3 Vpu virus showed an intermediate phenotype, with some stabilisation of p-p105 observed, while the profiles of cells infected with S3/7A Vpu and Δ Vpu viruses resembled uninfected cells. Similar profiles for p-IκBα were observed in infected CD4+ T cells (**Supplementary Figure 2a**).

Results thus far show that levels of phosphorylated p105 are increased in cells infected with viruses expressing 2_87 Vpu, both at steady state and following TNFα treatment. Considering that p105 has a complex role in the NF-κB pathway, both as the precursor to p50 and as a non-classical IκB, with phosphorylation important for both processes, this could have important implications. We therefore sought to clarify the effect of Vpu on p105 in transient p105 processing assays. N-terminally HA-tagged p105 constructs were transfected alongside an NF-κB stimulus (IKKβ) in the presence or absence of Vpu (**Figure 4e**). An increase in p50 levels consistent with signal-induced processing can be seen in response to co-expression of IKKβ, along with the appearance of an upper p105 band indicative of phosphorylated or monoubiquitinated p105. In contrast to **Figures 4a-d**, however, the presence of 2_87 Vpu does not result in the stabilisation of p105; rather, the upper p105 band is no longer present, and p50 levels are depleted. p105 and p50 levels in the presence of S3/7A and NL4.3 are similar to stimulated p105 processing in the absence of Vpu. The same pattern was observed when using TNFα as a stimulus (**Supplementary Figure 2b**). Again, increased processing of p105 to p50 in the presence of active NF-κB signalling was significantly diminished in cells co-expressing 2_87 Vpu. To demonstrate that Vpu specifically affects processing of p105 to p50, rather than acting directly on the p50 protein, identical assays were performed using HA-tagged p50 rather than p105. Levels of p50 were maintained under all conditions (**Supplementary Figure 2b**). As Vpu has a marked effect on p105 and a downstream effect on p50, we performed p50 nuclear translocation assays as an alternative to more traditional p65 translocation assays. P105 constructs were N-terminally tagged with mCherry and upon co-transfection with IKKb, p105 processing led to p50 translocation to the nucleus (**Figure 4f**). As expected, the presence of 2_87 Vpu inhibited p50 translocation, while S3/7A and NL4.3 Vpus were unable to inhibit nuclear translocation, or in some cases demonstrated an intermediate phenotype (**Figure 4f**).

### Primary Vpu inhibits the non-canonical NF-κB pathway

The non-canonical equivalent of p105 is p100, which becomes phosphorylated by IKKα following stimulation and is partially proteasomally processed to p52 (**Figure 1b**). Analogous to p105, it also functions as an IκB and can assemble into high molecular weight complexes containing multiple NF-κB dimers (kappaBsomes) [42-45]. It has also been implicated in downstream signalling following activation of cytosolic DNA sensing pathways [46, 47]. P100 processing assays, conducted by co-transfecting N-terminally-tagged p100 with NIK, demonstrated that 2_87 Vpu was able to inhibit the processing of p100 to p52 (**Figure 5a**), while both S3/7A and NL4.3 were defective. Effects on both the non-canonical and canonical pathways, induced by AZD5582 and TNFα respectively, were next compared under conditions of natural infection, using NL4.3 viruses expressing either 2_87, 2_87 S3/7A or NL4.3 Vpu in HEK293T cells (**Figure 5b**) and CD4+ T cells (Jurkat, **Figure 5c**). Cells were infected for 42 hours then treated for 6 hours before harvest. In uninfected cells, the induction of the non-canonical pathway by AZD5582 was indicated by the detection of phospho-p100 and the increased processing of p100 to p52, while TNFα stimulation resulted in increased levels of p100 and the detection of phospho-p105. As previously shown in **Figure 4**, infection with the 2_87 Vpu virus resulted in increased levels of p-p105 in untreated cells, and this was much more pronounced upon treatment with TNFα; for the S3/7A and NL4.3 viruses, p-p105 levels were similar to uninfected cells upon treatment. Indicative of the interdependence of the two pathways [48], AZD5582 stimulation also lead to the stabilisation of p-p105, and this was higher in cells infected with the 2_87 Vpu virus. Under the same conditions, a strong and striking stabilisation of p-p100 is seen, accompanied by a reduction in p100 processing, represented by the reduction of p52 levels, and increase in p100 levels, back to those seen in uninfected, unstimulated lanes. P-p100 and p100/p52 levels in cells infected with S3/7A Vpu virus were similar to uninfected cells, while NL4.3 Vpu virus gave an intermediate phenotype. As also shown in **Figure 2**, β-TrCP was diminished in cells infected with 2_87-expressing virus. Overall, these results demonstrate that infection with viruses possessing optimal Vpu function causes significant dysregulation of both canonical and non-canonical NF-κB pathways.

**Figure 5.**
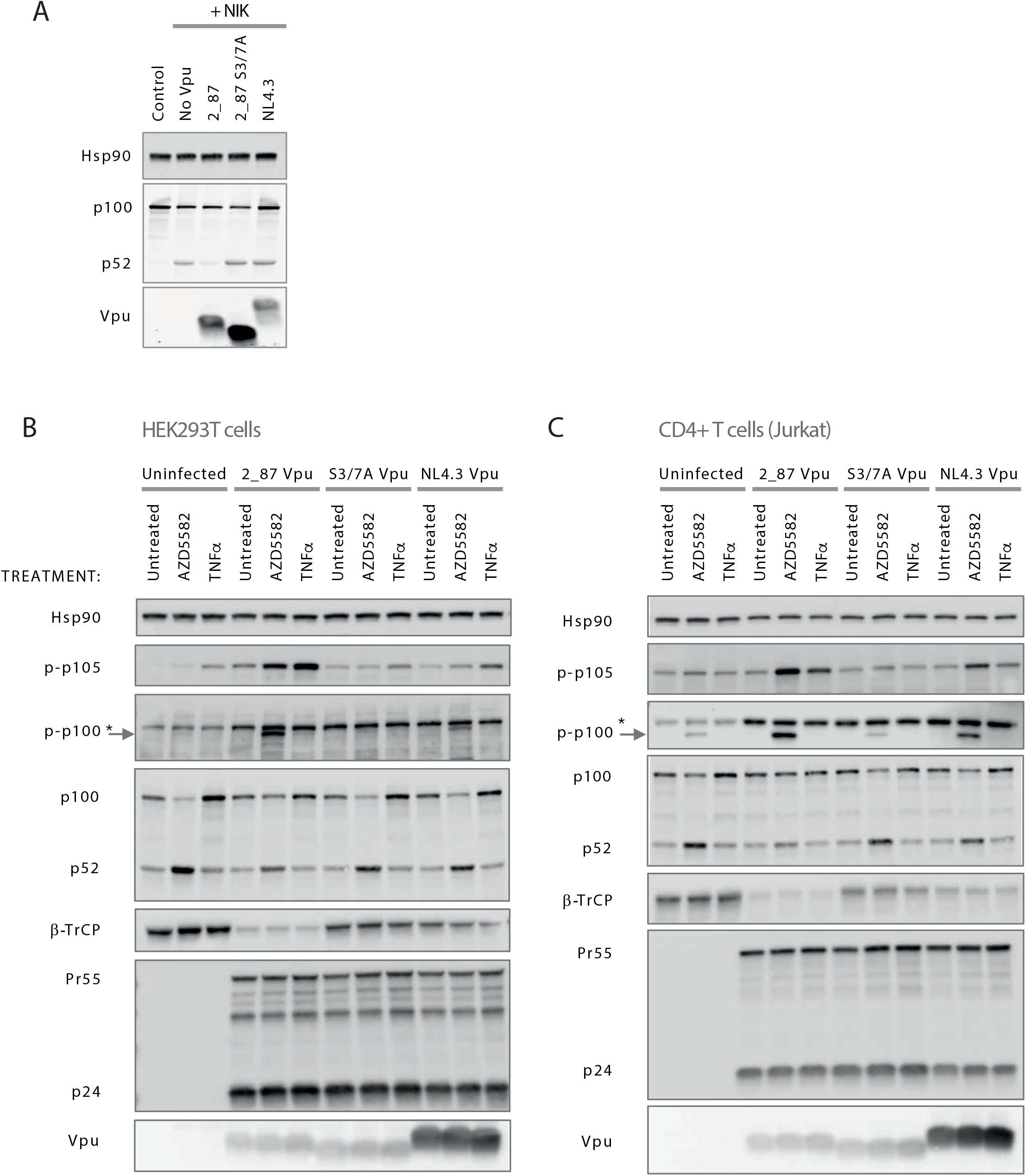
Vpu inhibits the processing of p100 to p52 and leads to the stabilisation of phospho-p100 in infected cells. **(A)** Transient p100 processing assays were performed by co-transfecting HA-p100, NIK and Vpu (2_87, 2_87 S3/7A or NL4.3) plasmids into HEK293T cells. 24 hours after transfection, cells were harvested and western blotted for HA (p100, p52 and Vpu) and Hsp90 as a loading control. **(B)** Recombinant NL4.3 proviruses engineered to express either 2_87, 2_87 S3/7A or NL4.3 Vpu were used to infect HEK293T cells at an MOI of 3. 42 hours after infection, cells were treated with 200 nM AZD5582 or 10 ng/ml TNFα, or left untreated. 6 hours after treatment, cells were harvested and western blotted for Hsp90 (loading control), phospho-p105 (Ser932), phospho-p100 (Ser866/870), total p100, p52 and β-TrCP. * denotes non-specific band. HIV-1 Gag (p55 and p24) and Vpu were blotted as controls for infection levels. (**C**) as for (**B**) but using CD4+ T cells (Jurkat).

### Both β-TrCP1 and -2 must be knocked down to phenocopy the effects of 2_87 Vpu

The reduction of β-TrCP1 and the stabilisation of β-TrCP2 by 2_87 Vpu prompted us to question whether there was a hierarchy in these actions for the inhibition of NF-κB by Vpu. As previously reported, Vpu specifically co-opts the SCF^β-TrCP^ to target tetherin for degradation in the host cell [33, 35]. In agreement with these studies, knocking down β-TrCP1 expression by siRNA had no effect on the ability of 2_87 and NL4.3 Vpus to down-regulate cell surface CD4 expression, whereas β-TrCP2 knockdown alone, or in combination with β-TrCP1, led to a restoration, albeit partial, of cell surface CD4 levels (**Figure 6a**). In contrast, individual knockdowns had a marginal effect on the canonical and non-canonical pathways as measured by p105 and p100 phosphorylation (**Figure 6b**), whereas the double knockdown of β-TrCP1 and -2 phenocopied the hallmarks of NF-κB inhibition by 2_87 Vpu, with the significant stabilisation of p-p105 and p-p100, indicating that the inhibition of both β-TrCP paralogues is required for the inhibition of NF-κB by Vpu.

**Figure 6.**
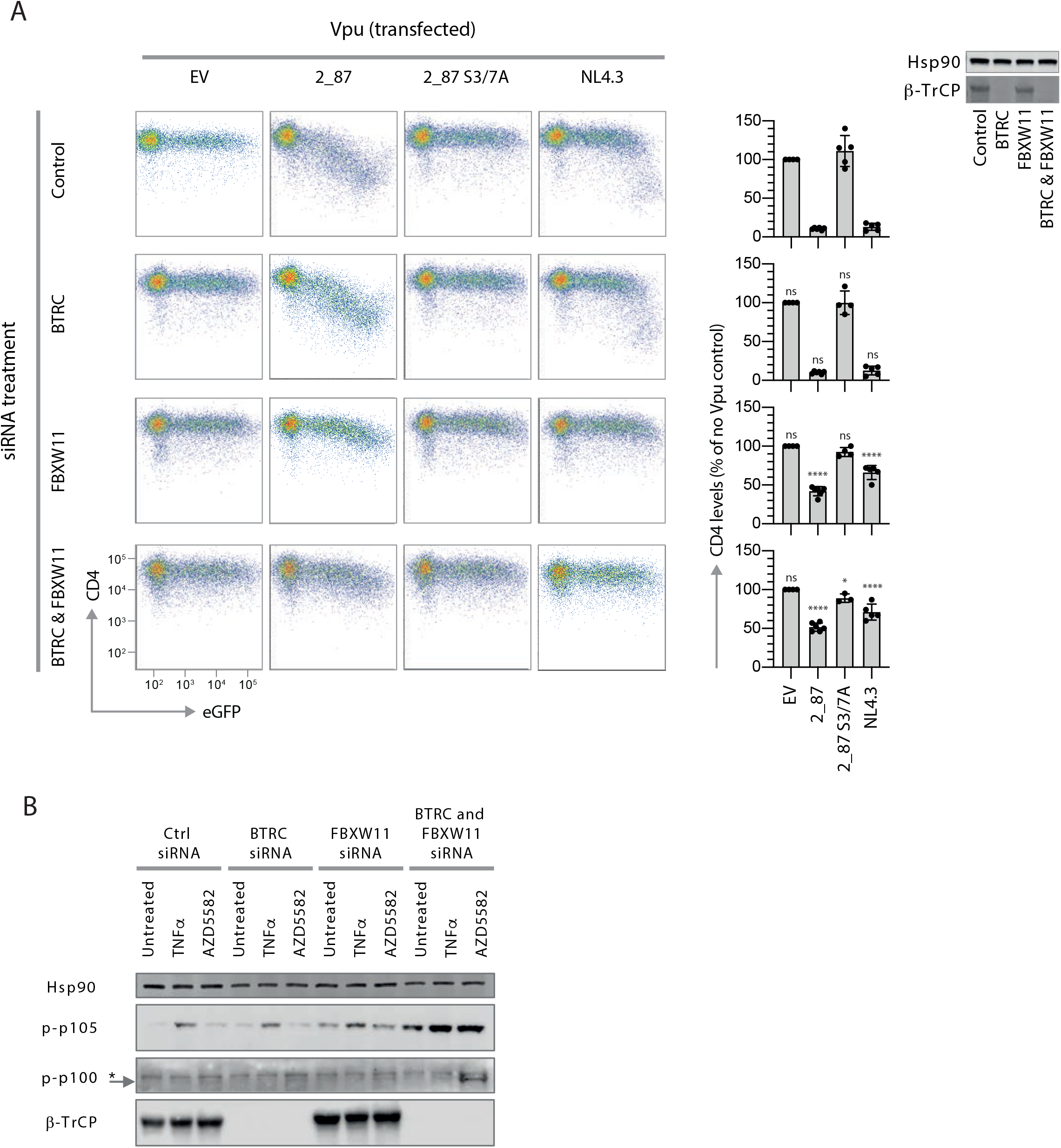
siRNA knockdown of β-TrCP2 is required to inhibit CD4 cell-surface downregulation by Vpu, but knockdown of both paralogues is required to phenocopy 2_87 Vpu NF-κB inhibition. **(A)** Prior to CD4 downregulation assays, CD4+ TZMbl cells were pre-treated with siRNA to downregulate β-TrCP1 (BTRC), β-TrCP2 (FBXW11) or both. CD4 downregulation assays were performed by co-transfecting Vpu and GFP, harvesting 24 hours later and analysing cell surface CD4 expression of gated GFP-positive cells by flow cytometry. Results are normalised to CD4 median fluorescent intensity in the absence of Vpu (EV). Graphs show means from at least 4 independent experiments ± SD. Asterisks above the bars indicate significant differences seen for each siRNA treatment compared to untreated cells, calculated separately for each Vpu: *p* value >0.1 (ns), <0.1 (*), <0.01 (**), <0.001 (***), <0.0001 (****). β-TrCP1 levels in siRNA-treated cells are shown by western blot, with Hsp90 as loading control. **(B)** HEK293T cells were pre-treated with siRNA for β-TrCP1 (BTRC), -2 (FBXW11) or both, then treated with 10 ng/ml TNFα, 200 nM AZD5582 or left untreated for 6 hours before harvesting. Lysates were analysed by western blot for Hsp90 (loading control), phospho-p105 (Ser932), phospho-p100 (Ser866/870) and β-TrCP.

### Determinants of Vpu required for binding and degradation of β-TrCP1

We next focused on features of Vpu that contribute to the inhibition of NF-κB activity, in particular the contribution of individual serine residues. Following initial reports demonstrating that both serines in the SGNES motif are phosphorylated [26, 27, 49], S52/56 or S53/57 have traditionally been mutated together, and their contribution to Vpu function is rarely investigated individually, particularly in the context of primary Vpu. Therefore, all three serines in the cytoplasmic tail of 2_87 Vpu (**Figure 7a**) were individually mutated to alanines. We found that mutating serine 57 had a greater effect on NF-κB inhibitory activity than mutating serine 53, and that the S57A mutant closely resembled wildtype NL4.3 in its inhibitory profile (**Figure 7b**). As reported previously for other functions of Vpu, mutating serine 65 enhanced the NF-κB inhibitory function of Vpu [50]. In contrast to the 2_87 profiles, mutating either serine 52 or 56 in NL4.3 had a similar negative impact on function with no dominant effect of either serine (**Figure 7b**).

**Figure 7.**
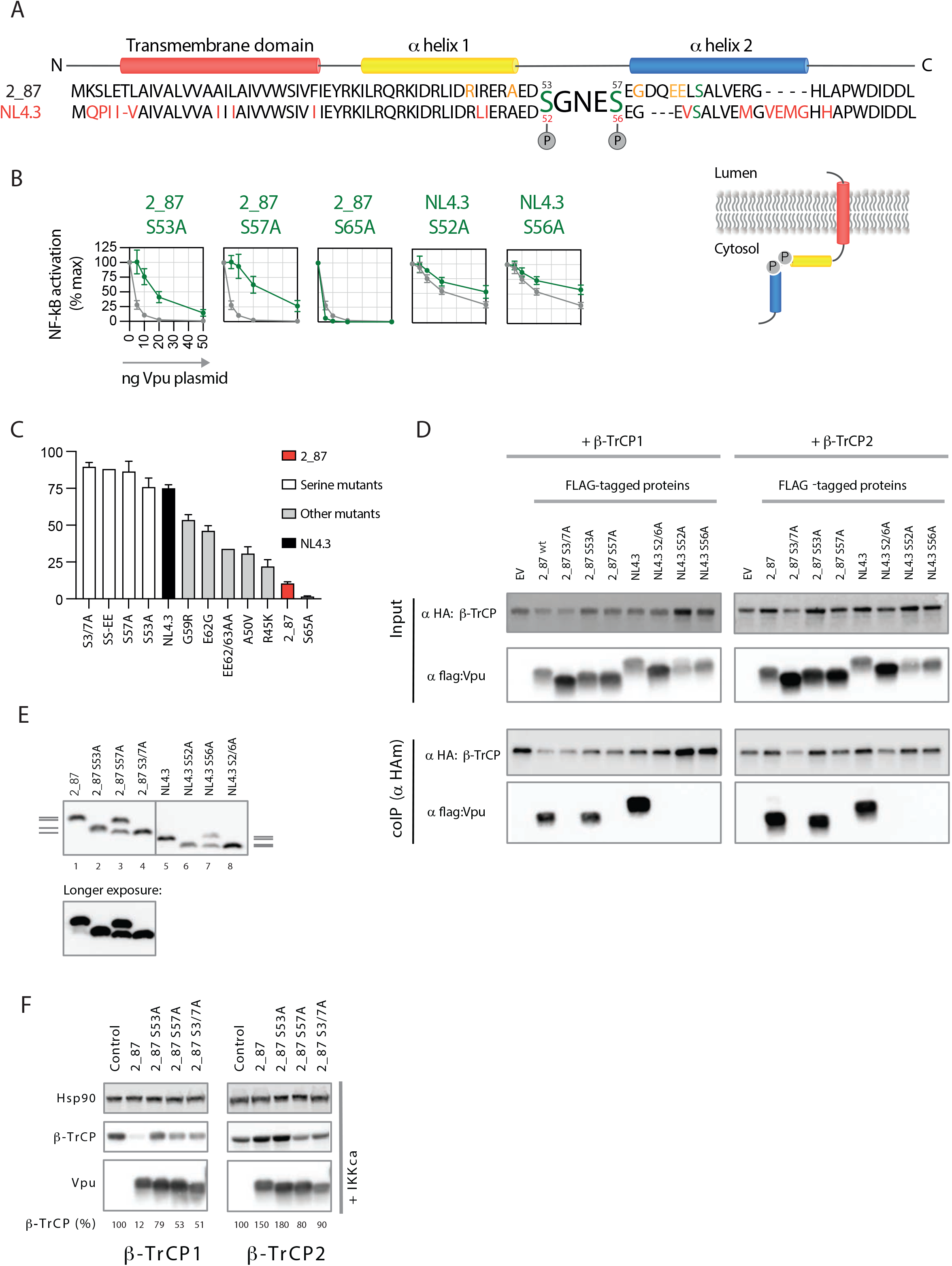
For 2_87 Vpu, serine 53 is sufficient for binding to β-TrCP whereas NL4.3 Vpu requires both serines. Both 2_87 serines are required for degradation of β-TrCP1. **(A)** Alignment of 2_87 and NL4.3 Vpu with domains indicated. Cytoplasmic tail serines are denoted in green. Residues in NL4.3 that differ from 2_87 are coloured red. Residues in 2_87 found to affect NF-κB inhibition in a screen of primary Vpus [31], and tested in panel **(C)** are shown in orange. **(B)** Transient NF-κB activation assays, using MAVS as a stimulus, were performed as for **Figure 1c**. Results are expressed as a percentage of normalised signal recorded in the absence of Vpu (% max). Means are presented from at least 3 independent experiments, with error bars showing ± SD. 2_87 single serine mutants are shown in green with 2_87 in grey on each graph, and NL4.3 single serine mutants are shown in yellow with NL4.3 in grey on each graph. **(C)** A panel of Vpus, including serine and combined serine mutants and naturally-occurring mutations that specifically impacted NF-κB inhibition [31] were compared for their ability to inhibit NF-κB induced by MAVS in transient NF-κB reporter assays. Mutants are arranged in order of impact. Wildtype 2_87 is shown in red. NL4.3 is shown in black. Serine mutants are shown in white. Mutations found to impact NF-κB inhibition in a primary Vpu screen and made in the 2_87 Vpu background are shown in grey and depicted in (**A**). **(D)** Single serine mutants of 2_87 (S53A and S57A) and NL4.3 (S52A and S56A) Vpus were compared for their ability to bind β-TrCP1 and -2 by immunoprecipitation. Dual serine mutants of each Vpu (2_87 S3/7A and NL4.3 2/6A) were used as negative controls. HEK293T cells were co-transfected with flag-tagged Vpu or EV and HA-tagged β-TrCP or EV, and 24 hours later cells were lysed, immunoprecipitated with anti-HA antibody and analysed by western blot. **(E)** HEK293T cells were transfected with HA-tagged 2_87 or NL4.3 Vpu and single- and double-serine mutants thereof. Cell lysates were resolved by phosphate-affinity PAGE, on 10% polyacrylamide gels containing 50uM Phos-tag™. Western blots were probed with anti-HA antibody to demonstrate the phosphorylation states of 2_87 and NL4.3 Vpus and corresponding single- and dual-serine mutants. Grey lines on the side of the gels indicate defined phosphorylation states for 2_87 Vpu (left) and NL4.3 Vpu (right). **(F)** The direct effect of individual serine mutants of Vpu on β-TrCP1 and -2 was examined by co-transfecting HEK293T cells with HA-tagged β-TrCP1 or -2 plus Vpu, in the presence (+IKK) of active signalling. 24 hours after transfection, cell lysates were harvested and analysed by western blot for HA (β-TrCP and Vpu) and Hsp90 (loading control).

A previous study in which we compared the ability of 304 primary Vpus to counteract physical virus restriction by tetherin with the inhibition of tetherin-mediated NF-κB signalling revealed regions of Vpu that were specifically required for the counteraction of NF-κB activation. All residues were located in regions flanking the SGNES site. These naturally-occurring mutations were introduced into a 2_87 Vpu background and tested for their ability to inhibit MAVS-stimulated NF-κB. All mutants were compared at a dose of 10ng, at which input 2_87 Vpu reduces NF-κB activation induced by MAVS by 90% (**Figure 1c** and **Figure 7c**). As shown in **Figure 7c**, individually the mutants had a partial impact on NF-κB inhibitory function but none more so than the serine mutations. Mutants that affected the charge of an acidic patch C-terminal to the SGNES, G59R and E62G, had the greatest effect on function, potentially through disrupting the putative function of this region as a CK2 priming site [51]. Mutating the double aspartic acid residues to alanine in this acidic patch (EE62/63AA) demonstrated a similar phenotype.

In order to determine whether differences in β-TrCP binding accounted for the results shown in **Figure 7b** and **c**, co-IP experiments were performed with the single serine mutant Vpus. Mutating serine 57 completely abolished binding of 2_87 Vpu to both β-TrCP1 and -2, in keeping with the more potent effect of this mutation on Vpu function, whereas mutating serine 53 had no effect on binding (**Figure 7d**). This result suggests hierarchical serine phosphorylation, which is consistent with CK2 requiring an acidic amino acid (aspartic or glutamic acid) or a phosphorylated serine/threonine at position +3 of the phosphorylation site (minimal consensus sequence [S/T]xx[E/D/S^p^];[51]), and agrees with original reports of preferential CK2 phosphorylation of serine 56 of NL4.3 Vpu *in vitro* [26]. Interestingly, the co-IP experiments revealed another prominent difference between the NL4.3 and 2_87 Vpus: while the presence of serine 57 was sufficient for the binding of 2_87 Vpu to β-TrCP1 and -2, mutation of either of the serine residues in NL4.3 Vpu completely abolished binding to both β-TrCP paralogues (**Figure 7d**).

We reasoned that disrupted phosphorylation of NL4.3 Vpu compared to 2_87, on either of the serines but in particular S56, might account for the requirement for both serines for binding to β-TrCP and for the resemblance of the 2_87 S57A NF-κB inhibitory profile to that of NL4.3 (**Figure 7b**). To investigate this, we performed phosphate-affinity PAGE, which specifically resolves phosphorylated proteins (**Figure 7e**). In agreement with previous studies [52], the profile for 2_87 Vpu showed four clearly defined phosphorylation states: double phosphorylation (lane 1), single phosphorylation on S57 (lane 2), single phosphorylation on S53 plus unphosphorylated (lane 3), and unphosphorylated (lane 4). Consistent with the results from co-IP experiments and with sequential phosphorylation, the phosphorylation state of the S57A mutant was partial, whereas the phosphorylation of the S53A mutant, although a smaller gel shift, was total. NL4.3 showed a similar profile, with the caveat that bands were less resolved for NL4.3 than for 2_87. Thus, serine phosphorylation differences are unlikely to account for the reduced potency of NF-κB inhibition by NL4.3 Vpu or the total inhibition of binding to β-TrCP upon mutation of either serine; it is more likely that differences in the regions flanking the SGNES contribute to the defect.

Finally, the effects of the individual 2_87 serine mutants were further explored in direct β-TrCP co-transfection assays (**Figure 7f**). Individual serine mutants had an impaired impact on β-TrCP1 levels (**Figure 7f**, left panel), whereas S53A maintained the ability to stabilise β-TrCP2 levels (**Figure 7f**, right panel). Thus, despite different impacts on β-TrCP binding, both serines are required for β-TrCP1 depletion by 2_87 Vpu.

## Discussion

Viruses have evolved to maintain an optimal balance between exploiting cellular processes for virus replication while inhibiting those that cause obstructions. This balance is exemplified by HIV requiring NF-κB for viral transcription, while also employing multiple strategies to temporally inhibit NF-κB signalling to avoid the induction of inflammatory responses at specific stages in its lifecycle. It is further illustrated by the co-opting of SCF^β-TrCP2^ by HIV-1 Vpu for the ubiquitination and subsequent degradation of its cellular targets, while inducing the degradation of β-TrCP1 to inhibit NF-κB. By inserting a potent primary Vpu (2_87) and mutants thereof into the NL4.3 provirus/GFP system we have demonstrated that Vpu both degrades and sequesters β-TrCP, and through these interactions has multi-layered consequences for the infected cell.

Beyond HIV-1 Vpu, other proteins from diverse viral families contain decoy degrons that bind and disable β-TrCP, including rotavirus NSP-1 (DSGIS; [53-58]), EBV LMP1 (DSGHES; [59]) and vaccinia virus A49 (YSGNLES; [37]), all of which are mechanistically distinct. Following phosphorylation of its C-terminal degron by CKII, NSP-1 binds to β-TrCP and recruits a Cul3 CRL complex via its N-terminal RING domain, triggering the poly-ubiquitination and degradation of β-TrCP [53-58]. Thus, NSP1 acts as the CRL substrate adaptor while β-TrCP becomes the substrate. A49, on the other hand, is not associated with β-TrCP degradation but acts as a transdominant decoy, binding and sequestering β-TrCP via its phospho-serine motif [37]. A49 employs a further regulatory step through the phosphorylation of its decoy degron by the IKK complex itself, thus only being activated upon triggering of the signalling cascade it subsequently inhibits [38]. For EBV Lmp1, certain variants of this oncogenic protein demonstrate a biphasic activation of NF-κB – activating at moderate levels then inhibiting at high expression levels, due to dominant negative inhibition of β-TrCP through the LMP1 decoy degron [59]. As demonstrated here and previously, for HIV-1 Vpu, the binding of the SGNES degron to β-TrCP has dual functionality: to act as a dominant negative decoy molecule, binding β-TrCP and preventing its participation in the NF-κB pathway [28-30], and to recruit the SCF^β-TrCP^ ligase for the ubiquitination and subsequent degradation of its target cellular proteins, including CD4 and tetherin [23, 32-36]. We further demonstrate that this involves inducing the degradation of the β-TrCP1 paralogue in a CRL-dependent manner, while exploiting β-TrCP2 for the recruitment of the Cul1 CRL complex. Thus, while evolving an SxxxS motif might seem a simple act of molecular mimicry adopted by multiple virus families to inhibit the NF-κB pathway, the regulation and specific mechanisms behind such inhibitory strategies are varied and complex.

To date, Vpu remains one of the few substrates of β-TrCP that distinguishes between the two paralogues. In agreement with previous studies demonstrating that degradation of tetherin requires only β-TrCP2 [33, 35], but in conflict with the requirement for both β-TrCP1 and β-TrCP2 for the degradation of CD4 [32], we show that only β-TrCP2 is required for the downregulation of CD4. Despite undetectable levels of β-TrCP1 in siRNA experiments, both 2_87 and NL4.3 Vpus were able to achieve full downregulation of CD4 from the cell surface (**Figure 6**). This is logical given the significant depletion of β-TrCP1 seen in cells infected with virus expressing the 2_87 Vpu (**Figure 2** and **Figure 5b**,**c**) and the disparity between β-TrCP1 and -2 levels in the presence of 2_87 Vpu in transient assays (**Figure 3a**,**c** and **Figure 7f**). Thus, for the ubiquitination and degradation of Vpu’s cellular targets, as for rotavirus NSP1, Vpu becomes the substrate adaptor, connecting the SCF^β-TrCP2^ ligase machinery to CD4 and tetherin, without being degraded itself. For the β-TrCP1 depletion, it remains to be determined whether this involves an active degradation similar to that seen with NSP1, with β-TrCP1 becoming the substrate, or whether Vpu exploits a feature of β-TrCP1 regulation and turnover that differs from β-TrCP2. Of note, we have excluded the possibility that Vpu co-opts a β-TrCP2-specific SCF CRL to degrade β-TrCP, as there was no discernible difference in β-TrCP levels in cells treated with β-TrCP2 siRNA (**Supplementary Figure 1c**).

The simultaneous inhibition and co-option of β-TrCP by Vpu has multi-layered consequences for the infected cell, including the inhibition of both the canonical and non-canonical NF-κB pathways, in addition to effects on other myriad targets of β-TrCP such as CDC25A and β-catenin [16, 60-62]. In cells infected with viruses expressing primary Vpu proteins we demonstrate potent stabilisation of both phosphorylated p100 and p105. These proteins have a complex role in the NF-κB pathway, acting both as IκBs (IκBδ and γ respectively) and precursors to mature NF-κB subunits p52 and p50. As IκBs, they are able to form high-molecular weight complexes – in the case of p100 these are known as kappaBsomes − that sequester multiple NF-κB subunits, estimated to inhibit up to 50% of the cellular NF-κB [44, 45]. Thus, their inhibition likely accounts for the increased potency of NF-κB inhibition by primary 2_87 Vpu, and indeed, perhaps that of other viral proteins that target β-TrCP.

Whether the inhibition of the non-canonical pathway is directly beneficial for HIV-1, or whether it is simply a side effect of inhibiting the canonical pathway, requires further investigation. Non-canonical signalling has been demonstrated to occur in a cGAS/STING-dependent pathway following the sensing of cytoplasmic DNA [46, 47], therefore the targeting of this pathway by Vpu may serve to further evade undesirable signalling events in the infected cell. Furthermore, promising latency reversal strategies that use SMAC mimetics to activate the noncanonical NF-κB pathway and re-activate integrated viral genome transcription, while avoiding the more pleiotropic effects of canonical NF-κB agonists such as PKC activators [63, 64], would need to take into account the potent inhibitory effects of primary HIV-1 Vpu proteins on this pathway demonstrated here. Humanised mouse experiments using the cell line-adapted JR-CSF Vpu, and macaque experiments using SIVmac, however, may underestimate the counteractive effect of primary Vpus on this pathway.

As previously noted by us and Sauter et al. [11], the binding of Vpu to β-TrCP does not strictly track with the ability of Vpu to inhibit NF-κB. Primary Vpus with double serine mutations have residual NF-κB inhibitory function, despite being unable to bind β-TrCP [11, 31]. We extend those findings by demonstrating that only S57 is essential for binding to β-TrCP1 and -2, yet this mutant had NF-κB inhibitory activity equivalent to that of NL4.3 Vpu. Interestingly, despite no apparent differences in phosphorylation status (**Figure 7e**), NL4.3 is significantly diminished for NF-κB inhibitory activity. Single serine mutations of either serine 52 or 56 completely abolishes binding of NL4.3 Vpu to β-TrCP, yet the overall ability of wildtype NL4.3 to bind β-TrCP1 and -2 does not appear impaired in co-immunoprecipitation or immunofluorescent experiments (**Figure 3b,c** and **Figure 7d**). The obvious region of Vpu to account for such differences is the second alpha helix of the cytoplasmic tail, where there are multiple differences between 2_87 and NL4.3 Vpu, including the lack of additional acidic residues in NL4.3, predicted to act as a CKII prime site. Indeed, some of the natural mutations found to specifically impact NF-κB inhibition mapped to this region [31].

Early studies demonstrated constitutive phosphorylation of Vpu [26, 27, 49]. Here we demonstrate clear differences between NL4.3 and 2_87 Vpus in the requirement for phosphorylation on one or both serines for the binding of β-TrCP, with 2_87 Vpu only requiring the phosphorylation of S57. We further demonstrate that, while only S57 is required for 2_87 Vpu binding to β-TrCP1 and -2 and the stabilisation of β-TrCP2, both serines are required for degradation of β-TrCP1 (**Figure 7f**). Proteomics studies have indicated that Vpu can potentially interact with [65] and be dephosphorylated by [66] the phosphatase PP2A. Furthermore, an additional serine at position 61 (65 in 2_87 Vpu) has been shown to regulate Vpu function and lead to its proteasomal degradation via an unidentified CRL [50]. The phosphorylation state of Vpu has also been predicted to determine its oligomerisation status [52]. Thus, further studies are required to understand the precise phosphorylation status of all three serines in the Vpu cytoplasmic tail in infected cells, and how this may relate to the regulation of its myriad functions.

We have demonstrated that infection with HIV-1 leaves behind a tell-tale trace of NF-κB perturbation, in the form of phosphorylated p105, even in the absence of exogenous stimuli (**Figure 4** and **Figure 5b,c**). In all infection conditions, including those in the absence of Vpu or presence of suboptimal Vpus, p-p105 could be detected. This rose to significantly higher levels of p-p105 in the presence of the highly active 2_87 Vpu. No such indication of signal activation was found in phospho-IκBα or p65 after 48 hours of viral infection, suggestive of long-term rather than acute activation of the pathway. It is unclear what provides the initial stimulus for this activation. Potentially some level of viral sensing may be at play, or this may reflect the activity of viral proteins shown to boost NF-κB signalling such as gp41 [67] or Nef [11]. Thus, while the detection of phospho-p105 reveals the NF-κB activation that occurs due to HIV infection and is exploited for viral transcription, the inhibition of the pathway mediated by Vpu is in turn apparent in the significant enrichment of p-p105. As such, p-p105 has potential as a convenient marker for NF-κB status in infected cells.

In summary, we provide a detailed view of the consequences of β-TrCP inhibition in the HIV-1-infected cell, including previously undocumented interference with the non-canonical NF-κB pathway. We underscore the importance of using Vpu proteins representative of natural infection and studied in the context of actively replicating virus.

## Materials & methods

### Cells

HEK293T cells and Jurkat T cells were obtained from American Type Culture Collection (ATCC). HeLa TZMbl were obtained through the NIH HIV Reagent Program, Division of AIDS, NIAID, NIH, kindly provided by John C. Kappes. Primary CD4+ T cells were purified from freshly isolated PBMCs from healthy donors. PBMCs were isolated by density gradient using Lymphoprep (Axis-Shield) and CD4+ T cells purified by negative selection using the Dynabeads Untouched Human CD4+ T Cell Isolation kit (Invitrogen) according to the manufacturer’s instructions. CD4+ T cells were activated using Human T-Activator CD3/CD28 Dynabeads (Invitrogen) according to the manufacturer’s instructions and maintained in RPMI GlutaMax supplemented with 10% FCS and 30 U/ml recombinant IL-2 (Roche).

### Ethics

Ethical approval to draw blood from healthy donors as a source for primary lymphocytes was granted by the KCL Infectious Disease BioBank Local Research Ethics Committee – approvals SN1-100818 and SN1-160322

### Western blot analyses

Cell lysates were resolved on gradient gels (4-8%; BioRad) and blotted onto nitrocellulose membranes. Unless otherwise stated, all blots were incubated at 4°C overnight in 5% BSA, using the following antibodies: mouse anti-HA antibody (anti-HA.11 clone 16B12, BioLegend UK Ltd.); rabbit anti-HA antibody (#600-401-384, Rockland Inc.); rabbit anti-Flag antibody (#F7425, Sigma-Aldrich, UK); mouse and rabbit anti-Hsp90 (Santa Cruz). β-TrCP (D13F10) rabbit mAb (#4394); IKKβ (D30C6) rabbit mAb (#8943); phospho-IKKα (Ser176)/IKKβ (Ser177) (C84E11) rabbit mAb (#2078; blocked with SuperBlock (Thermo Scientific)); IκBα rabbit mAb (#9242); phospho-IkBa (Ser32/36) (5A5) mouse mAb (#9246); NF-κB1 p105/p50 (D4P4D) rabbit mAb (#13586); phospho-NF-κB p105 (Ser932) (18E6) rabbit mAb (#4806); NF-kB2 p100/p52 (D7A9K) rabbit mAb (#37359); phospho-NF-κB2 p100 (Ser866/870) rabbit mAb (#4810; blocked with SuperBlock (Thermo Scientific)); NF-κB p65 (D14E12) rabbit mAb (#8242); phospho-NF-κB p65 (Ser536) (7F1) mouse mAb (#3036; blocked with 5% milk); all from Cell Signaling Technology. Anti-HIV-1 p24 mouse mAb (183-H12-5C) was kindly provided by Dr Bruce Chesebro and Kathy Wehrly through the NIH HIV Reagent Program, Division of AIDS, NIAID, NIH (#ARP-3537). Anti-HIV-1 Vpu rabbit antibody was kindly provided by Andrés Finzi [9, 68].

### Phosphate affinity PAGE

10% polyacrylamide gels were prepared containing 50uM PhosTag (Alpha Laboratories) in the separating gel. Cell lysates were first diluted 1:10 in llaemmli buffer, and SDS-PAGE was performed using standard protocols using methanol-based transfer.

### Plasmids

pCR3.1 myc-β-TrCP2/FBXW11 has been described previously [41]. Constructs with N-terminal GFP and HA tags were made by subcloning β-TrCP2 into pCR3.1 GFP and HA. Human β-TrCP1/BTRC was cloned into pCR3.1 myc, HA and GFP for the expression of N-terminally tagged β-TrCP1. Human IKKβ was cloned into pCR3.1 for the expression of C-terminally FLAG-tagged protein. A constitutively active version (SS177,181EE) was generated by quick-change site-directed mutagenesis using Phusion-II polymerase (New England Biolabs) and standard protocols. The pCR3.1 tetherin plasmid has been previously described [39]. The MAVS expression plasmid was kindly provided by Jeremy Luban. 3xκB-pConA-FLuc and pCMV-RLuc renilla control were kindly provided by Andrew Macdonald [69]. Human p105/NFKB1 was cloned into pCR3.1 HA and CHE for the expression of HA- and mCherry-N-terminally tagged p105, resulting in tagged expression of both the full-length unprocessed p105 and processed p50. A truncated version was generated for the expression of N-terminally HA-tagged p50 only. Likewise, human p100/NFKB2 was cloned into pCR3.1 HA for the expression of N-terminally tagged p100 and tagged processed p52. Human NIK was cloned into pCR3.1. pCR3.1-Vpu-HA plasmids expressing C-terminally tagged codon-optimised Vpus NL4.3, NL4.3 double serine mutant SS52/56AA (“S2/6A”), 2_87 and 2_87 double serine mutant SS53/57AA (“S3/7A”) have been described previously [41, 70]. Flag-tagged equivalents were generated by subcloning. Mutants were generated by quick-change site-directed mutagenesis (NL4.3 S52A and S56A; 2_87 S53A, S57A, S65A, R45K, A50V, G59R, E62G, EE62/63AA and the 2_87 double serine phospho-mimetic SE53/57EE or “SS-EE”) using Phusion-II polymerase (New England Biolabs) and standard protocols. The A49 expression plasmid was kindly provided by Geoffrey Smith [37].

### Proviral constructs

An HIV-1 proviral construct (HIV-1 NL4-3 IRES-eGFP infectious molecular clone that encodes the full length HIV-1 NL4.3 genome with the *nef* open reading frame augmented by an IRES-eGFP (kindly provided Drs Munch, Schindler and Kirchhoff via the NIH HIV Reagent Program, Division of AIDS, NIAID, NIH (pBR43leG-nef+, cat #11349 [71])) was used as the basis of all viruses described. This proviral genome was rendered Vpu-defective as described previously [70]. The 2_87 Vpu was inserted in place the NL4.3 Vpu by overlapping PCR to ensure sequences immediately upstream of the Vpu start codon known to be important for Vpu and Env expression were unchanged [72]. Site directed mutations of the serine codons to alanines at positions 52/53 and 56/57 in NL4.3 and 2_87 Vpu were performed by quick-change. All viral genomes were sequence verified prior to use.

### Virus production

Sub-confluent HEK293T cells in 10 cm plates were co-transfected with 10 µg of proviral plasmid and 2 µg of pCMV-VSV-G plasmid using 1 mg/ml polyethyleneimine (PEI). Media was changed 6-12 hours post transfection. Cell supernatant was harvested 48h after transfection, filtered and ultracentrifuged over 20% sucrose in PBS at 28,000 rpm for 2 hours. Pellets were resuspended in serum-free RPMI medium, aliquoted and stored at -80°C. Titres (infectious units/mL) were determined on HeLa-TZMbl reporter cells.

### Transient NF-κB reporter assays

Reporter constructs expressing firefly luciferase under the control of an NF-κB promoter (3xκB-pConA-FLuc ([69]) were used for transient NF-κB inhibition assays. As detailed previously [31, 39], sub-confluent HEK293T cells were co-transfected in 24-well plates with 20ng 3xκB-pConA-FLuc, 10ng pCMV-RLuc renilla luciferase control plasmid, stimulus plasmid (10 ng pCR3.1 MAVS/IPS1/Cardif, 50ng pCR3.1 tetherin/BST2, 20ng pCR3.1-IKKβ-flag, or 20ng pCR3.1 NIK HA) or equivalent quantity of empty vector control, and 10 ng of pCR3.1 Vpu HA or empty vector control. In the case of titration experiments, 5, 10, 20, 50 and 100 ng of pCR3.1 Vpu HA plasmid were used and supplemented to 100ng with empty vector plasmid. 24 hours after transfection, cells were harvested and both firefly and renilla luciferase activity measured with the Dual-Luciferase Reporter Assay System (Promega), according to the manufacturer’s instructions. Firefly luciferase was normalised to the renilla signal, and fold NF-κB activation for each stimulus calculated relative to empty vector control in the absence of Vpu expression.

### Transient β−TrCP degradation assays

Cells were transfected with 120 ng pCR3.1-HA-β-TrCP1 or 50 ng pCR3.1-HA-β-TrCP2 plasmid plus 25 ng of pCR3.1-IKKβ-flag or empty vector, plus 50 ng of pCR3.1 Vpu HA plasmid (2-87, NL4.3 or mutants thereof) or empty vector per well of a 24 well plate. 24 hours after transfection cell lysates were harvested for Western blot analyses.

### P105 or p100 processing assays

For p105 processing assays, sub-confluent HEK293T cells plated in 24w plates were co-transfected with 100ng pCR3.1-HA-p105 plus 20ng pCR3.1-IKKβ-flag or empty vector plus 50ng pCR3.1 Vpu HA or empty vector and harvested 24 hours later for western blot analyses. In the case of TNFα stimulation, cells were transfected as above but omitting IKKβ flag, and treated with 10 ng/mL TNFα 18 hours after transfection. Cells were harvested at 5, 15, 30, 60, 120, 240 and 360 minutes after TNFα addition and analysed by western blot. For p100 processing assays, cells were co-transfected with 100ng pCR3.1-HA-p100 plus 100ng pCR3.1 NIK or empty vector plus 50ng pCR3.1 Vpu HA or empty vector and harvested 24 hours later for western blot analyses.

### siRNA knockdown assays

Cells were pre-treated with siRNA prior to CD4 downregulation assays, TNFα or AZD5582 treatment or infection. ON-TARGETplus SMARTpool human BTRC siRNA and human FBXW11 siRNA (Dharmacon) were used to target β-TrCP1 and -2 respectively. TZMbl or HEK293T cells were reverse transfected with 20 pmol siRNA per well of a 24 well plate using Lipofectamine RNAiMAX transfection reagent (ThermoFisher Scientific) according to the manufacturer’s instruction. 24 hours later, cells were trypsinised, split into 3 and the reverse transfection process was repeated. 24 hours after the second reverse transfection, cells were transfected, treated with TNFα or AZD5582 or infected.

### CD4 downregulation

Cells were pre-treated with siRNA prior to CD4 downregulation assays, as detailed above. CD4 downregulation assays were performed as described previously [31]. Briefly, 24 hours after the second siRNA treatment, sub-confluent TZMbl cells were co-transfected with 200 ng pCR3.1-GFP or empty vector control and 40 ng pCR3.1 Vpu or empty vector control in 24 well plates. 24 hours after transfection, cells were harvested and stained for cell surface CD4 expression using anti-human CD4 APC (RPA-T4, eBioscience, ThermoFisher Scientific) and analysed by flow cytometry on a BD FACSCanto II system (BD Biosciences) using FlowJo software. Cells were gated for high GFP expression and CD4 levels were determined as median fluorescence intensity in the absence of Vpu expression, with CD4 levels in the presence of Vpu expressed as a percentage of this.

### TNFα and AZD5582 NF-κB activation assays

HEK293T or Jurkat CD4+ T cells were infected as above and treated with TNFα or AZD5582 for 6 hours before harvest at 48 hours post infection. For TNFα timecourse assays, cells were infected in bulk at an MOI of 3, then divided into separate wells (300,000 cells per well) for TNFα or control treatment. TNFα was added 42 hours after infection, to a final concentration of 10 ng/ml, and cells were harvested at 0, 15, 30, 60, 120 and 240 minutes post-TNFα addition for western blot analyses.

### Virus infection assays

HEK293T cells were infected in 24-well plates at 300,000 cells per well at an MOI of 3 or 5. Jurkat and CD4+ T cells were infected in 48-well plates at 500,000 cells per well at an MOI of 3 or 5. 48 hours after infection cells were harvested for western blot analyses. For drug treatment of infected cells, a final concentration of 10µM Mg132, 100µM MLN4924, 50nM concanamycin A or mock treatment of DMSO or water as appropriate was added to the cell culture medium 6 hours before harvest.

### Immunofluorescence

For p50 nuclear translocation assays, sub-confluent HEK293T cells plated in 24w plates were co-transfected with 100 ng pCR3.1 CHE p105, 25 ng pCR3.1-IKKβ-flag or empty vector, and 20 ng pCR3.1 Vpu HA (2_87, 2_87 S3/7A or NL4.3) or empty vector. For β-TrCP localisation assays cells were co-transfected with 150 ng of pCR3.1 GFP BTRC or 100 ng of pCR3.1 GFP FBXW11 and 20 ng of pCR3.1 Vpu HA (2_87, 2_87 S3/7A or NL4.3) or empty vector. For microscopy, glass coverslips were placed in 24-well plates and treated with 400ul of 10% gelatin in PBS (pre-warmed at 37°C to liquify) for 30 minutes at room temperature. To obtain optimal cell density for microscopy of individual cells, 24 hours after transfection each well was trypsinised, split 1:4 to 1:7 and re-seeded onto the pre-treated glass coverslips. Remaining cells were replated into 24w plates for parallel western blot analysis as required. Cells were allowed to adhere to the glass cover slips overnight at 37°C, then fixed with 4% formaldehyde in PBS for 15 minutes at room temperature, washed once with PBS then with 10mM glycine in PBS. To permeabilise, cells were treated with 0.1% Triton X-100 and 1% BSA in PBS for 15 minutes at room temperature, before incubation with mouse anti-HA antibody (anti-HA.11 clone 16B12, BioLegend) in 0.01% Triton X-100 in PBS for 45 minutes at room temperature. Cells were washed three times with 0.01% Triton X-100 in PBS, incubated with Alexa Fluor 488, 594 or 647 anti-mouse secondary antibody (Molecular Probes, Invitrogen) and washed again three times. Cover slips were mounted on slides with ProLong™ Diamond Antifade Mountant with DAPI (Invitrogen) and imaged on a Nikon Eclipse Ti inverted microscope with Yokogawa CSU-X1 spinning disk unit. Image analyses were performed with NIS Elements Viewer and Fiji software.

### Immunoprecipitation

Subconfluent HEK293T cells were co-transfected with 600 ng pCR3.1-β-TrCP1/BTRC-, β-TrCP2/FBXW11-HA or empty vector control, plus 500 ng pCR3.1 Vpu flag or empty vector control per well of a 6-well plate. 26-28 hours after transfection, cells were lysed in IP buffer (50 mM Tris pH 7.5, 100 mM NaCl, 1 mM EDTA, 2mM DTT, 0.1% Nonidet P40 substitute, supplemented with cOmplete™ Protease Inhibitor Cocktail (Roche) and PhosSTOP™ phosphatase inhibitor tablets (Roche)), incubated for 10 mins on ice, sonicated and centrifuged for 5 mins at 13,000 rpm at 4°C. Lysates were incubated with mouse anti-HA antibody (anti-HA.11 clone 16B12, BioLegend) or rabbit anti-Flag antibody (F7425, Sigma) for 1 hour at 4°C with rotation. 60 ul of washed protein G agarose beads were added to each sample and incubated at 4°C with rotation overnight. Beads were then washed with IP buffer and resuspended in Laemmli buffer for western blot analysis. In the case of β-TrCP1 (BTRC)/Vpu CoIPs, cells were treated with 10 µM MG132 for 6 hours before harvest, to avoid degradation of β-TrCP1 by Vpu.

### Statistics

Statistical analyses were performed in Graphpad Prism v 9. Unless stated otherwise, all graphs show means from at least 3 independent experiments with errors bars indicating ± SD. Transient NF-κB reporter assays and CD4 downregulation assays with siRNA treatment were analysed using two-way ANOVA with multiple comparisons and mixed-effects analyses. Western blot intensities were calculated by first normalising to the Hsp90 loading control for each lane, and calculating the percentage band intensity relative to the uninfected control. For phosphorylated targets, bands were further normalised within each gel to a positive control band containing stimulated, uninfected cell lysate (p-IκBα, p-p105 or p-p100). Normalised values from at least three independent experiments were then compared using unpaired one-tailed T tests (p-p105) or unpaired two-tailed T tests (β-TrCP). *p* value >0.1 (ns), <0.1 (*), <0.01 (**), <0.001 (***), <0.0001 (****).

## Acknowledgements

We are grateful to members of the Neil lab, past and present, for helpful discussion and technical assistance, in particular Helin Sertkaya, Irina Jahin and Sudeep Bhushal. We thank Andrés Finzi for kindly gifting the anti-Vpu antibody, Andrew Macdonald for reporter plasmids, Jeremy Luban for the MAVS expression plasmid and Geoffrey Smith for the A49 expression plasmid.

## Author Contributions

The study was conceived and designed by SP and SJDN. Experiments were performed by SP, JS and CK. Data were analysed by SP. The manuscript was written by SP and SJDN.

## Figure legends

**Supplementary Figure 1.**
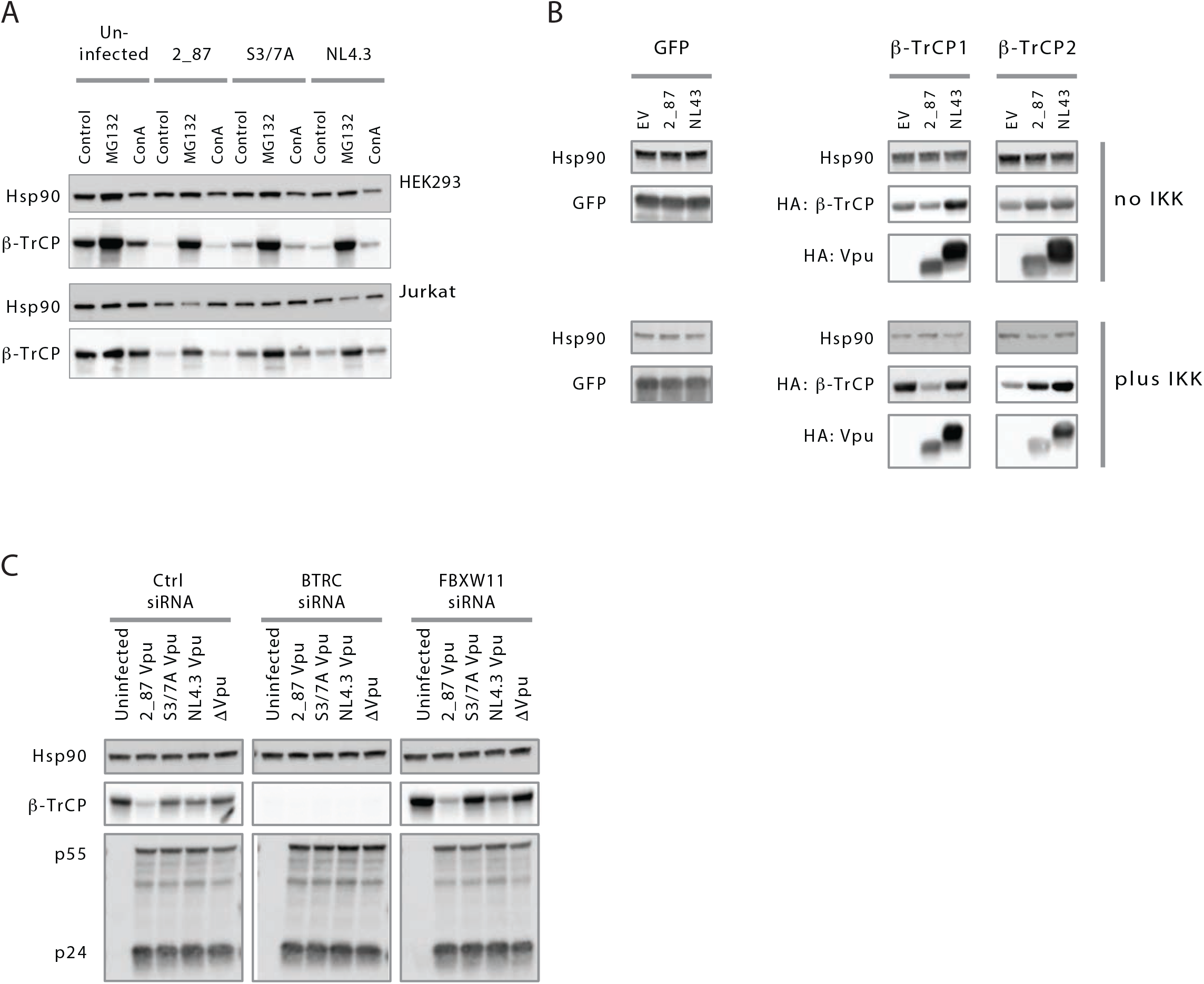
**(A)** HEK293T and Jurkat cells were infected with recombinant NL4.3 proviruses engineered to express either highly active 2_87 Vpu, 2_87 S3/7A Vpu, NL4.3 Vpu or no Vpu (Δ Vpu) at an MOI of 5, and treated with proteasomal inhibitor MG132 (10 µM) or concanamycin A (50 nM) for 6 hours prior to harvest at 48 hours. Cell lysates were analysed by western blot for Hsp90 (loading control) and endogenous β-TrCP levels. **(B)** HEK293T cells were co-transfected with GFP or HA-tagged β-TrCP1 or -2 plus Vpu or empty vector control, in the presence (+IKK) or absence (no IKK; empty vector) of active signalling. 24 hours after transfection, cell lysates were harvested and analysed by western blot for GFP, HA (Vpu and β-TrCP) and Hsp90 (loading control). **(C)** HEK293T cells were pre-treated with siRNA for β-TrCP1 (BTRC), -2 (FBXW11) or control before infecting with recombinant NL4.3 proviruses engineered to express either highly active 2_87 Vpu, 2_87 S3/7A Vpu, NL4.3 Vpu or no Vpu (Δ Vpu) at an MOI of 4. 48 hours after infection cells were lysed and analysed by western blot for Hsp90 (loading control), endogenous β-TrCP levels and HIV-1 Gag (major bands show p55 and p24).

**Supplementary Figure 2.**
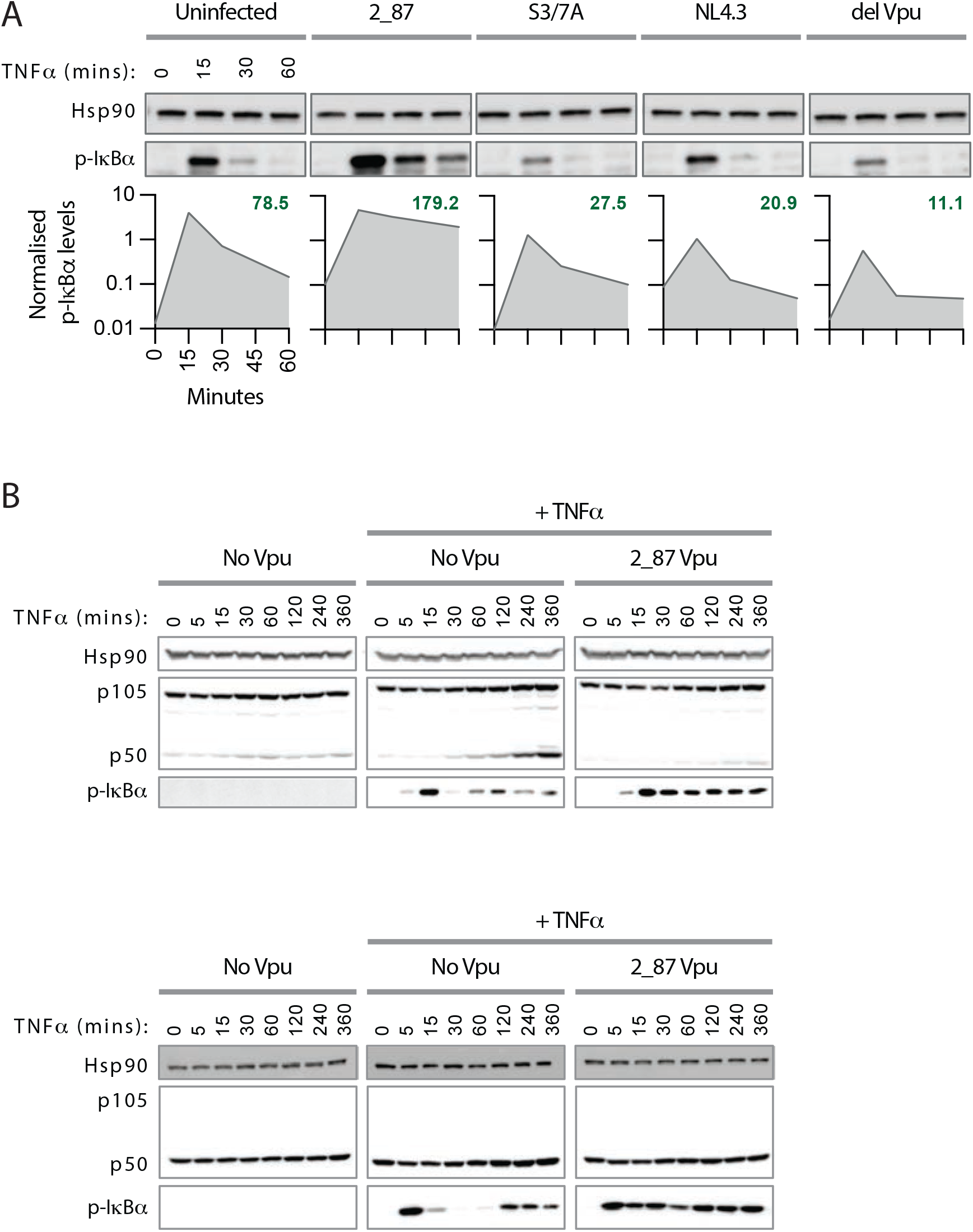
**(A)** IκBα stabilisation timecourse in Jurkat cells. CD4+ Jurkat T cells were infected with viruses expressing either 2_87, 2_87 S3/7A, NL4.3 or no Vpu (Δ Vpu) at an MOI of 3. 48 hours after infection, cells were treated with 10 ng/ml TNFα, and time points were harvested at 0, 15, 30 and 60 minutes following treatment. Samples were analysed by western blot for Hsp90 (loading control) and phospho-IκBα. Band intensities for p-IκBα are shown below each blot, normalised to Hsp90 for each sample and to positive controls for p-IκBα, as appropriate, per blot (not shown in the image). Numbers shown in bold green text on each graph represent the calculated area under the curve (AUC). **(B)** p105 processing timecourse with TNFα. Transient p105 processing assays were performed by co-transfecting HA-p105 (top panel) or HA-p50 (bottom panel) with 2_87 Vpu or empty vector control. 18 hours after transfection, cells were stimulated with 10ng/ml TNFα, or mock treated, and time points harvested for western blotting at 0, 5, 15, 30, 60, 120, 240 and 360 minutes. Blots were probed for HA (p105 and 50 in the top panel; p50 only in the bottom panel) and Hsp90 (loading control). Phospho-IκBα blots were included as positive controls for NF-κB signal activation.

